# Chronic neuropathic pain alters reversal learning and prefrontal cortex activation without generalized deficits in reward-related behavior

**DOI:** 10.64898/2026.02.24.707747

**Authors:** Andrea Aurelio Borges, Mitchell A. Nothem, Christina M. Curran-Alfaro, Karina Possa Abrahao, Jacqueline M. Barker

**Affiliations:** Department of Pharmacology and Physiology, Drexel University College of Medicine, Philadelphia, PA, USA; Departamento de Psicobiologia, Escola Paulista de Medicina, Universidade Federal de São Paulo, SP, Brazil; Departamento de Fisiologia, Escola Paulista de Medicina, Universidade Federal de São Paulo, SP, Brazil; Programa de Pós-Graduação em Psicobiologia, Escola Paulista de Medicina, Universidade Federal de São Paulo, SP, Brazil; Neuroscience Graduate Program, Drexel University College of Medicine, Philadelphia, PA, USA

**Keywords:** Chronic pain, sucrose self-administration, reversal learning, extinction

## Abstract

Chronic pain is associated with neuropsychiatric comorbidities characterized by impairments in cognitive and behavioral flexibility, yet the direct impact of chronic pain on reward-guided behavior remains poorly understood. Here, we investigated how chronic neuropathic pain altered sucrose-reinforced behavior across multiple behavioral assays with distinct cognitive-behavioral demands in male and female mice. Mice underwent spared nerve injury (SNI) or sham surgery and were tested in operant sucrose self-administration, reversal learning, and extinction training. The impact of acute painful stimulation on sucrose seeking was also assessed. SNI produced stable mechanical hypersensitivity without affecting acquisition or extinction of sucrose self-administration, indicating intact baseline acquisition and sucrose seeking. In contrast, chronic neuropathic pain selectively altered behavioral flexibility: SNI females exhibited faster acquisition of reversal learning, while SNI mice as a group showed impaired inhibition of responding on the previously reinforced lever during early reversal. Acute painful stimulation suppressed sucrose seeking in males, but not in females, independent of chronic pain status. At the neural level, painful mechanical stimulation differentially modulated medial prefrontal cortex (mPFC) activity, increasing infralimbic c-Fos expression in SNI mice while decreasing it in sham controls relative to non-stimulated animals. There was no evidence of chronic mPFC hyperactivity as indexed by ΔFosB expression. Together, these findings demonstrate that chronic neuropathic pain did not globally disrupt sucrose reward-related behavior, but instead selectively altered behavioral flexibility and pain–reward interactions in a sex-dependent manner, with accompanying alterations in mPFC engagement following aversive stimulation.

## Introduction

Chronic pain is defined as pain that persists for more than three months [66]. Chronic pain is maladaptive and is associated with risk for a range of neuropsychiatric disorders [1,39,49], including those defined by altered affect and behavioral flexibility [6,11,32]. Flexible reward seeking is impaired in substance use disorder [23,24] and depression [74], both of which have increased prevalence in people with chronic pain [5,21,36,58,69]. Neuropathic pain in particular – which results from disease or damage to the nervous system – is associated with depression-related outcomes in both clinical populations and in preclinical models [38,63,68]. However, the direct impact of chronic pain on reward-related behavior is not fully understood, with a range of findings in the preclinical literature. For example, recent work showed preserved cognitive performance in an experimental knee osteoarthritis rodent model of chronic pain [27], while others reported disrupted reward seeking and processing in chronic pain (c.f., [15]), including altered motivation for opioids [14,41], conditioned place preference [8], and reward seeking [31], though at least a subset of these effects may be specific to drug rewards [41]. Further, the impacts of chronic pain on reward and behavioral outcomes may be sex-dependent in preclinical models [12], consistent with sex differences in co-occurring chronic pain and neuropsychiatric illness in clinical populations. Together, there is a need for greater clarity on the relationship between chronic neuropathic pain and reward seeking, which necessitates use of preclinical models, for clear dissociation of directional effects of chronic pain on behavioral outcomes.

Preclinical models can also provide insight into the neural substrates that are differentially engaged during reward seeking during chronic neuropathic pain. The medial prefrontal cortex (mPFC), a key region involved in executive functions and adaptive behavior, has been implicated in chronic pain and neurobehavioral disorders [4,37,40,54,57,59,61]. The mPFC plays an important role in the regulation of behavioral flexibility, characterized by the capacity to modify behavior to adapt to a changing environment [28,33,45,56,67]. Subregions of the mPFC may have distinct roles in flexibility during reward seeking. The infralimbic cortex (IfL), for example, is essential for extinction learning and habits, while the prelimbic cortex (PrL) has opposing functions [10,26,29,55]. The anterior cingulate cortex (ACC) is necessary for behavioral flexibility by resolving competition between simultaneous signals and guiding appropriate action selection [13,60].

To characterize the impact of chronic neuropathic pain on a battery of reward-related behavior, the effect of spared nerve injury (SNI) on reversal learning, extinction, and behavioral response to acute painful stimulation was assessed in male and female mice. Further, the effect of acute painful stimulation on mPFC activation was determined using immediate early gene expression as a putative marker of cellular activity. The SNI model was selected as it mimics patient symptomatology, is highly replicable, and yields robust and long-lasting hypersensitivity, allowing for longer training and testing procedures [30]. Together, we report facilitated reversal learning in SNI-injured females, and suppressed sucrose seeking by painful stimulation in male mice, suggesting that the impact of chronic pain on reward seeking may depend on behavioral domain and sex.

## Methods

### Animals

Forty-one adult C57Bl/6J mice were used in this study (18 females and 23 males) from Jackson Laboratories. Mice were ordered at 9 weeks old and weighed 17-30 g at the beginning of the experiments. All animals were housed in a temperature-and humidity-controlled environment on a standard 12:12 light-dark cycle (07:00 on). Mice were handled and allowed to acclimate to the colony room for one week before beginning the behavioral experiments. Until the beginning of behavioral experiments, mice had *ad libitum* access to food and water. After surgery and during behavioral testing, animals were singly-house and food-restricted to maintain approximately 90% of their free-feeding body weight. All behavioral experiments were conducted between 10:00 AM and 12:00 PM. For group designations, mice were matched based on baseline paw withdrawal threshold (PWT). The Institutional Animal Care and Use Committee of Drexel University approved all procedures.

### Spared nerve injury surgery

Mice underwent spared nerve injury (SNI) or sham (as controls) surgery [20,53]. Animals were anesthetized with inhaled isoflurane, with induction at 4% and maintenance at 1–1.5%, and continuously monitored for respiration and general reflexes to ensure the proper depth of anesthesia was maintained. Body temperature was maintained using a heating pad throughout the procedure. Animals were shaved prior to surgery: An incision was made on the left hind leg, and the sciatic nerve was exposed. The common peroneal and tibial nerves were ligated with 6-0 silk suture and cut distally from the ligation. The skin was closed using size 7 wound clips (Reflex). The sural nerve was left intact. Sham mice underwent a similar procedure in which the branches of the sciatic nerve were exposed but not ligated or cut. Animals were allowed to recover for 2 weeks for the development of chronic hypersensitivity on the left paw before behavioral testing.

### von Frey assay

To assess allodynia, mice were stimulated on the lateral aspect of the plantar surface with a set of von Frey fibers using the up-down method [22] starting with a 0.16 g fiber (Stoelting Touch Test, Chicago, IL) and progressing to a 2.56 g fiber as the upper limit. PWT were calculated using the formula and appendix values described by Chaplan and colleagues [16] and employed in [53]. Animals were tested before surgery to obtain their baseline sensitivity, and at 3 different time points after surgery (Fig. 1A).

**Figure 1:**
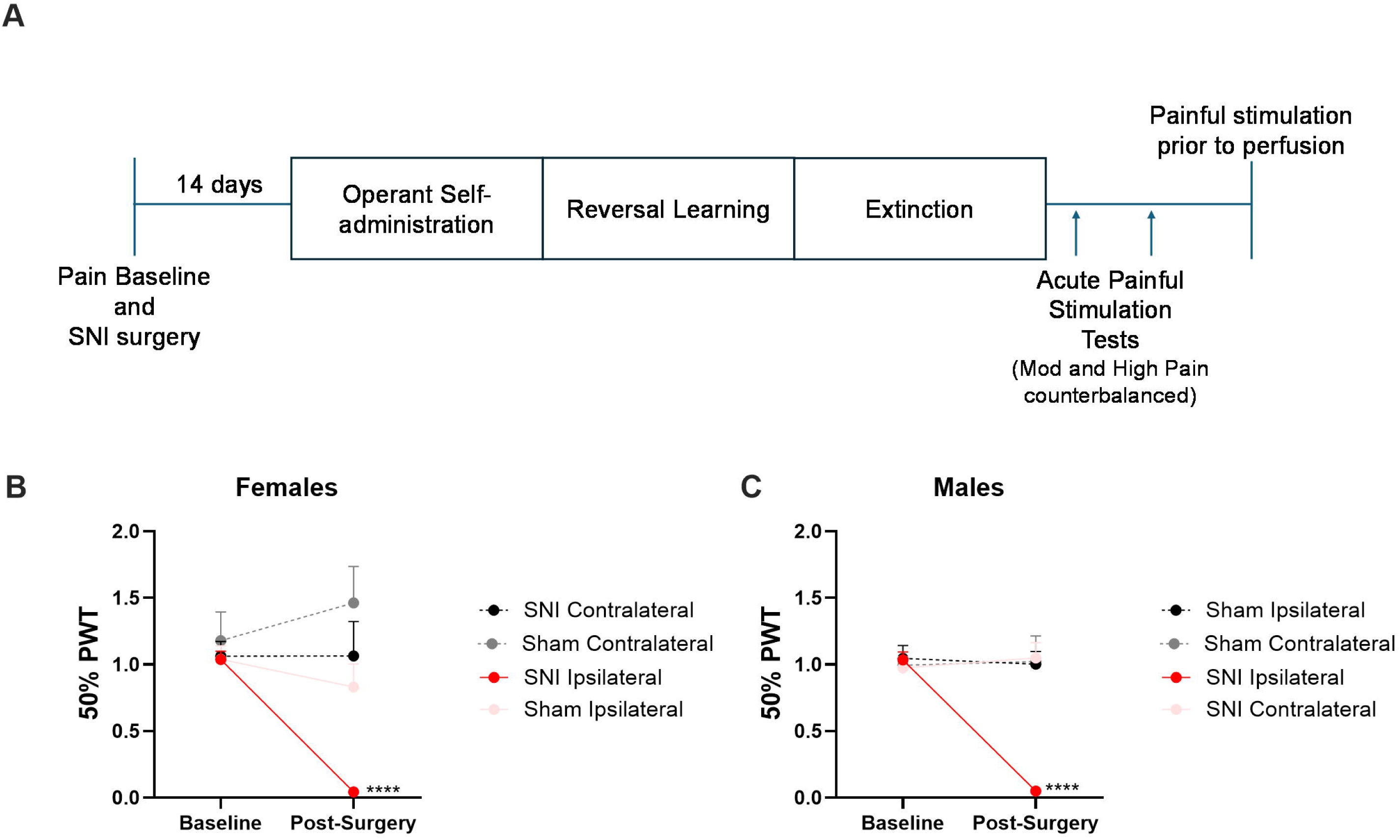
SNI surgery induced mechanical hypersensitivity in the injured paw. (**A**) Experimental timeline. (**B**) Paw withdrawal threshold (PWT) in the SNI females ipsilateral paw decreased significantly at 2 weeks compared with baseline and vs controls (n = 9/group). (**C**) PWT thresholds in the SNI males ipsilateral paw decreased significantly at 2 weeks compared with baseline and versus controls (n = 11/group). **** p < 0.001. Data are presented as mean ± SEM.

### Operant sucrose self-administration

#### Training phase

Two weeks after SNI or sham surgery, animals were food-restricted to approximately 90% of their original body weight. Mice were habituated to sucrose reward during 2 days in a standard operant conditioning chamber (21.59 x 18.08 x 12.7 cm; Med-Associates, Inc., Georgia, VT) where 20 uL of 10% sucrose solution was delivered in the magazine every 2 minutes on a fixed schedule with no levers available. During the acquisition phase, animals were trained daily to self-administer a 10% sucrose solution via lever pressing, with no cues indicating the delivery of reinforcement, requiring mice to acquire an action-outcome associative relationship between lever pressing and sucrose delivery. One lever delivered sucrose in the magazine (active lever), and the other did not (inactive lever). The reinforced levers were counterbalanced between groups. Daily training sessions lasted 30 minutes or until the animals completed 20 reinforced lever presses on a fixed ratio 1 schedule of reinforcement, where each lever press resulted in reward delivery. After 7 days, the schedule was shifted to a variable ratio 2 (VR2) for an additional 12 days. On this schedule, reward delivery occurred after a variable number of presses, averaging 2.

To determine whether self-administration was altered in SNI animals when reward delivery was not capped, the session was fixed at 30 minutes and total lever presses/reward deliveries were not capped for 3 additional consecutive training sessions. Reinforced lever presses, inactive lever presses, and magazine entries were recorded during all behavioral phases in the operant chamber. On the last day of the training phase, the cumulative number of reinforced lever presses was analyzed. “Magazine checking” behavior - defined as lever presses followed by a magazine entry - were also analyzed. Three animals (2 SNI males and 1 sham male) were excluded due to failure to acquire the task.

#### Reversal learning test

Reversal is a form of flexible behavior in which the active and inactive levers are switched, requiring the acquisition of a newly reinforced lever and/or the inhibition of responding on the previously reinforced lever. After completion of 3 30-min sessions, the reinforced lever was switched such that the previously inactive lever now delivered the sucrose reward, while the previously reinforced lever no longer did. The reinforcement schedule remained VR2 with one 30-minute session per day for at least 12 consecutive days. During initial reversal (the first two days of the reversal learning phase), magazine checking was also analyzed, defined as the percentage of lever presses followed by entry into the reward port, used to assess monitoring of the relationship between lever presses and reward delivery. Mice were considered to have reached the reversal criterion when they performed at least 70% of their lever presses on the newly reinforced lever across two consecutive sessions.

#### Extinction phase

To assess the ability to detect change in contingency when reward delivery was omitted, an extinction phase was implemented. During this phase, the reinforcer was withheld regardless of the animals’ responses. Sessions lasted 30 min per day for at least 5 consecutive days or until mice reached the extinction criterion, defined as fewer than 20% of lever presses compared to the average number of reinforced lever presses during the two days preceding extinction. One sham male failed to extinguish within 30-day sessions and was therefore excluded from further analysis.

#### Acute pain effects on sucrose seeking

To assess the effect of painful stimulation on sucrose seeking, animals underwent two seeking tests in extinction, following moderate and high pain stimulation, separated by additional extinction sessions. These tests were counterbalanced between cohorts to account for potential effects of repeated testing. On the moderate pain stimulation test day, animals underwent von Frey testing to determine the 50% PWT for each individual. Following PWT determination, animals received 10 stimulations on the lateral aspect of the plantar surface of the ipsilateral hindpaw relative to the injury, using a von Frey filament that delivered a force corresponding to approximately 50% of the PWT. The number of hindpaw responses was recorded. Stimulations occurred within 1–2 minutes. Immediately after stimulations, animals were placed in the operant chamber for a 30-minute session. Procedures for ‘moderate’ versus ‘high’ pain stimulation tests were identical, except for the force applied during the 10 stimulations. ‘Moderate’ stimulation tests used the von Frey filament closest to the calculated 50% PWT, while ‘high’ stimulation test employed two fibers above the 50% PWT, corresponding theoretically to approximately a 90% PWT. One SNI female was excluded from analyses due to low sensitivity.

### Brain tissue processing and immunohistochemistry

To assess mPFC activity regarding pain stimulation, expression of the immediate early gene c-Fos expression (marker of acute activity) and ΔFosB (marker of chronic activity) were assessed. Ninety minutes before perfusion, half of the animals were stimulated 10 times using a von Frey filament corresponding to a 90% PWT (high-intensity stimulated group). The remaining animals were placed on the testing platform for the same duration, but did not receive any stimulation (non-stimulated group). Mice were anesthetized with isoflurane and transcardially perfused with 1× PBS followed by 4% paraformaldehyde (PFA). Brains were post-fixed in 4% PFA for 24 h and cryoprotected in 20% sucrose for 24 - 48h. Coronal sections (40 µm) were cut on a cryostat (Leica GM 1510S), collected in quadruplicate, and stored in 0.01% sodium azide. Immunohistochemistry (IHC) was performed by incubating free-floating sections in 1% hydrogen peroxide for 1 h, followed by blocking in 5% normal donkey serum for 1 h. Sections were then incubated overnight at room temperature with primary antibodies against c-Fos (1:5000; 2250S, Cell Signaling) or ΔFosB (1:1000; 14695 S; Cell Signaling). The next day, sections were incubated with a biotinylated donkey anti-rabbit secondary antibody (1:1000; 711-065-152, Jackson Labs) for 30 min. Staining was visualized using ABC and nickel-enhanced DAB kits (PK6200 and SK-4100, respectively, Vector Labs) for 20 min. Sections were mounted on Superfrost Plus slides and coverslipped with DPX mounting medium. Images were acquired at 10x magnification in a Nikon Eclipse microscope with a panda SCMOS camera and stitched using Microsoft Image Composite Editor. ImageJ was used for automatic counting with a threshold-based approach in the anterior cingulate cortex (ACC), prelimbic cortex (PrL), and infralimbic cortex (IfL) in approximately three different slices (1.53 mm, 1.69 mm, and 1.97 mm anterior to bregma).

### Statistics

All statistical analyses and graphs were performed and designed using GraphPad Prism (version 10.6.1). Potential outliers were screened using Grubbs’ test, treating each data point as a potential outlier and evaluating all values individually. Analyses included two-way and three-way repeated-measures (rm) ANOVA as well as one-sample or two-sample t tests, as specified in the Results section. When sphericity was violated, the Geisser-Greenhouse correction was applied, and corrected degrees of freedom were reported. A mixed-effects model fitted with restricted maximum likelihood (REML) was employed for repeated-measures designs with missing values. Normality of the data was assessed using the D’Agostino-Pearson omnibus normality test. For all analyses, the threshold for statistical significance was set at α = 0.05. Following significant interactions, Sidák’s post hoc comparisons were performed.

## Results

### SNI surgery induced mechanical hypersensitivity in the injured paw

To confirm that mice developed abnormal sensitivity to mechanical stimulation after SNI, basal PWT on the ipsilateral and contralateral hindpaws was measured before and 2 weeks after SNI surgery using von Frey fibers. A rmANOVA on PWT in female mice revealed significant main effects of time [F_(1,32)_ = 5.33, p = 0.03], injury [F_(1,32)_ = 5.13, p = 0.03], and paw [F_(1,32)_ = 9.96, p = 0.004] and interactions between time × paw [F_(1,32)_ = 13.96, p = 0.0007], time × injury [F_(1,32)_ = 7.20, p = 0.01] and time × paw × injury [F_(1,32)_ = 1.62, p = 0.21]. Sidák’s post hoc test showed that PWT in the ipsilateral paws of SNI females were significantly reduced at 2 weeks compared to the corresponding baseline values (p < 0.0001). PWT did not change in the SNI contralateral paw or either paw in sham mice compared to their corresponding baseline values (all p’s > 0.50) (Fig. 1B). PWT in males were consistent with changes observed in females – a rmANOVA revealed significant main effects of time [F_(1,40)_ = 10.76, p = 0.002], injury [F_(1,40)_ = 10.49, p = 0.002] and paw [F_(1,40)_ = 9.43, p = 0.004] and interactions between time × paw [F_(1,40)_ = 16.90, p = 0.0002], time × injury [F_(1,40)_ = 10.05, p = 0.0003], injury × paw [F_(1,40)_ = 10.84, p = 0.002] and time × paw × injury [F_(1,40)_ = 12.31, p = 0.001]. Sidák’s post hoc test showed that PWT in the ipsilateral paw of SNI males were significantly reduced at 2 weeks compared to the corresponding baseline values (p < 0.0001). No changes were observed for PWT in the SNI contralateral paw or in either paw for sham animals (all p’s > 0.97) (Fig. 1C).

### SNI did not affect acquisition of operant chamber sucrose self-administration

Magazine entries during the 2 days of habituation to sucrose reward delivery in the operant chamber were analyzed. There were no significant effects of session [F_(1, 36)_ = 3.50; p = 0.07] or injury [F_(1, 36)_ = 0.003; p = 0.96] but there was a significant main effect of sex [F_(1, 36)_ = 10.00; p = 0.003] such that females made more magazine entries than males. There were no significant interactions between session × sex [F_(1, 36)_ = 0.55; p = 0.47], session × injury [F_(1, 36)_ = 0.008; p = 0.93], sex × injury [F_(1, 36)_ = 1.89; p = 0.18], or session × injury × sex [F_(1, 36)_ = 0.39; p = 0.54].

To determine if SNI impacted acquisition of sucrose self-administration, session length during the first 18 FR1 and VR2 sessions was assessed. These sessions were capped by number of reinforcers earned and thus shorter sessions reflect faster self-administration. A rmANOVA indicated that there was a main effect of session number on the session duration [F_(3.14,100.3)_ = 57.50, p < 0.0001, Geisser-Greenhouse corrected ε = 0.18], consistent with a reduction in the length of session required to reach the maximum of 20 correct lever presses decreased over time in all groups. There were no effects of sex [F_(1.32)_ = 0.29, p = 0.59] or injury [F_(1.32)_ = 1.91, p = 0.18], or interactions between session × injury F_(17, 544)_ = 0.45; p = 0.97], sex × injury [F_(1, 32)_ = 0.91; p = 0.35], or session × injury × sex [F_(17, 544)_ = 0.34; p = 0.99]. However, a sex x injury interaction was observed [F_(17,544)_ = 1.81, p = 0.02]. Corrected post hoc comparisons indicate that session length did not significantly differ between males and females on any session (all p’s > 0.05). In males, session length was reduced compared to session 1 on sessions 5-18 (all p’s < 0.05). In females, session length was not reduced compared to session 1 until session 7 (all p’s < 0.05) (Fig. 2A).

**Figure 2.**
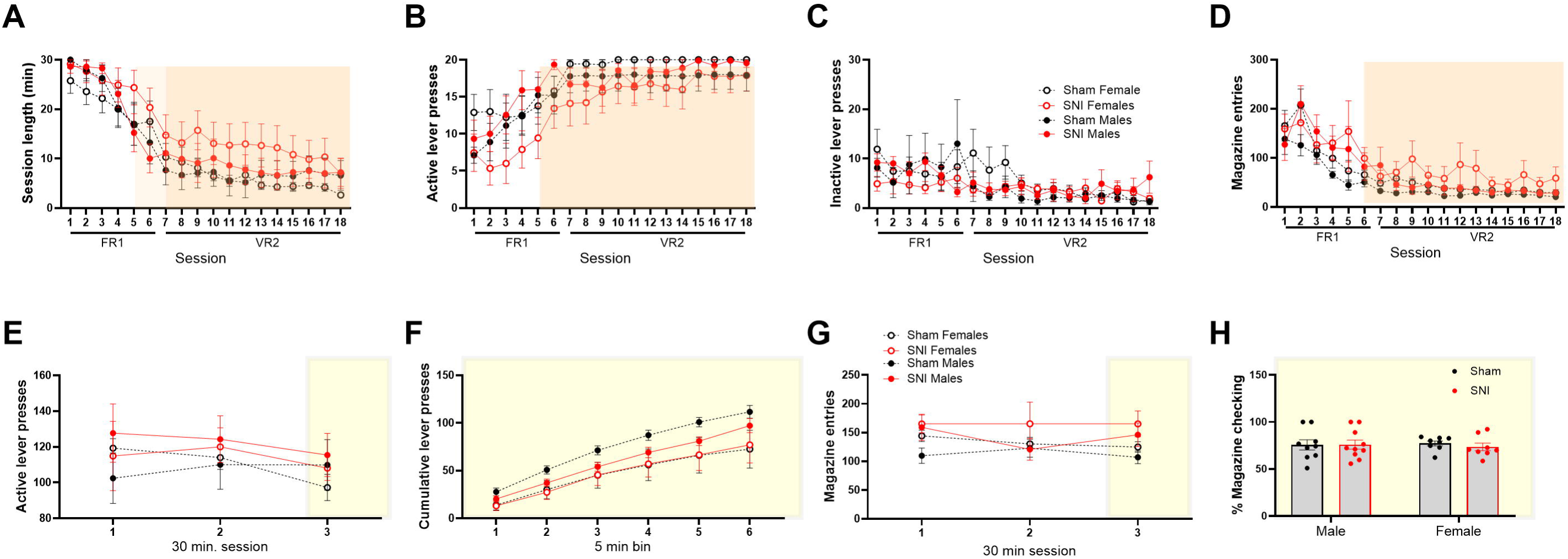
SNI does not affect acquisition of operant sucrose self-administration. (A) Session duration across the first 18 acquisition sessions (FR1 and VR2 schedules), which were capped at 20 reinforcers. Session length progressively decreased across training in all groups, reflecting acquisition of the task, with effects of injury. (B) Number of active lever presses across acquisition sessions increased over time in all experimental groups, with no effects of sex or injury. (C) Inactive lever presses decreased across sessions, indicating improved discrimination between active and inactive levers, with no effects of sex or injury. (D) Magazine entries during acquisition decreased across sessions in all groups, consistent with more efficient task performance, and were not affected by sex or injury. (E) Number of active lever presses during three subsequent fixed-duration (30 min) sessions following acquisition, showing no effects of session, sex, or injury. (F) Cumulative active lever presses across 5-min bins during the final 30-min session (yellow shadow) increased over time similarly in all groups, with no effects of injury. (G) Magazine entries during the fixed-duration sessions were not affected by session, sex, or injury. (H) Percentage of active lever presses followed by a magazine entry on the final training day (yellow shadow), showing no differences between sham and SNI mice or between sexes. Data are presented as mean ± SEM. n = 9–10 mice per group. Sham and SNI mice of both sexes acquired and performed sucrose self-administration similarly, indicating that SNI-induced hypersensitivity does not alter acquisition or baseline performance of operant sucrose seeking.

Consistent with this reduced session duration, a rmANOVA indicated that the number of active lever presses increased across sessions [F_(3.43,109.7)_ = 24.77, p < 0.0001, Geisser-Greenhouse corrected ε = 0.20]. Post hoc analyses indicated that active lever presses were greater on days 5-18 compared to day 1 (all p’s < 0.01; Dunnett’s multiple comparisons test). There were no effects of sex [F_(1,32)_ = 0.14, p = 0.71] or injury [F_(1,32)_ = 0.91, p = 0.35], no significant interaction between session × sex [F_(17, 544)_ = 1.43; p = 0.13] session × injury F_(17, 544)_ = 0.80; p = 0.70], sex × injury [F_(1, 32)_ = 2.45; p = 0.13], or session × injury × sex F_(17, 544)_ = 0.50; p = 0.95] (Fig. 2B).

The number of inactive lever presses decreased across sessions [F_(3.65, 113.1)_ = 4.81, p = 0.002, Geisser-Greenhouse corrected ε = 0.21] in all experimental groups (Fig. 2C). Post hoc comparisons indicated that inactive lever presses were reduced on sessions 8 and 10-18 compared to session 1 (all p’s < 0.05). There were no effects of sex [F_(1.31)_ = 0.002, p = 0.97] or injury [F_(1.31)_ = 0.19, p = 0.367] nor interactions between session × sex [F_(17, 527)_ = 0.97; p = 0.49] session × injury [F_(17, 527)_ = 1.364; p = 0.15], sex × injury [F_(1, 31)_ = 0.57; p = 0.46], or session × injury × sex [F_(17, 527)_ = 1.01; p = 0.46] were observed (Fig. 2C). Magazine entries also decreased across sessions [F_(4.73,165.4)_ = 31.49, p < 0.0001, Geisser-Greenhouse corrected ε = 0.28]. Post hoc comparisons indicated that magazine entries were lower on sessions 6 - 18 compared to session 1 (all p’s < 0.01). There were no effects of sex [F_(1.34)_ = 1.60, p = 0.21] or injury [F_(1.34)_ = 2.69, p = 0.11] nor interactions between session × sex [F_(4.72, 160.6)_ = 0.60; p = 0.70; Geisser-Greenhouse corrected ε = 0.28], session × injury F_(4.72, 160.6)_ = 1.13; p = 0.28, Geisser-Greenhouse corrected ε = 0.28], sex × injury [F_(1, 34)_ = 0.0004; p = 0.98], or session × injury × sex F_(4.72, 160.6)_ = 1,43; p = 0.22; Geisser-Greenhouse corrected ε = 0.28] (Fig. 2D).

After initial acquisition sessions, which were capped at 20 reinforcers, 3 sessions were performed where duration was fixed at 30 minutes, regardless of the number of lever presses. This was performed to assess whether self-administration was altered in SNI animals. Despite the increased time duration, there were no significant effects of session [F_(1.66, 56.27)_ = 2.75, p = 0.08, Geisser-Greenhouse corrected ε =0.83], sex [F_(1.24)_ = 0.05, p = 0.82], or injury [F_(1,34)_ = 0.69, p = 0.41] or interactions between session × sex [F_(1.66, 56.27)_ = 1.06, p = 0.34, Geisser-Greenhouse corrected ε = 0.83], session × injury [F_(1.66, 56.27)_ = 0.04, p = 0.94, Geisser-Greenhouse corrected ε = 0.83], sex × injury [F_(1, 34)_ = 0.22, p = 0.64], and session × sex × injury [F_(1.66, 56.27)_ = 2.01, p = 0.15, Geisser-Greenhouse corrected ε =0.83], consistent with similar self-administration patterns in sham-injured and SNI mice (Fig. 2E).

To determine if the pattern of active lever presses were different within session, cumulative responding was assessed across 5 minute bins in the final session. A rmANOVA indicated an effect of bin [F_(1.199,38.36)_ = 103.6; p < 0.0001, Geisser-Greenhouse corrected ε =0.24], with lever presses increasing significantly over time in all groups (Fig. 2E). Post hoc analysis indicated that responding was greater in all bins than bin 1 (all p’s < 0.001). No effects of sex [F_(1.32)_ = 3.88, p = 0.06] or injury [F_(1,32)_ = 0.52, p = 0.47] were observed, nor were there interactions between bin × sex [F_(5,160)_ = 1.72, p = 0.13], bin × injury [F_(5, 160)_ = 0.19, p = 0.96], sex × injury [F_(1.32)_ = 0.60, p = 0.44], or bin × sex × injury [F_(5,160)_ = 0.28, p = 0.92] on within session responding, consistent with similar sucrose self-administration in sham and SNI mice (Fig. 2F).

For magazine entries, there were no effects of session [F_(1.502, 51.05)_ = 0.68, p = 0.47, Geisser-Greenhouse corrected, ε =0.75], sex [F_(1.34)_ = 1.92, p = 0.18], or injury [F_(1,34)_ = 3.80, p = 0.18], or interactions of session × sex [F_(1.502, 51.05)_ = 0.09, p = 0.38, Geisser-Greenhouse corrected, ε =0.75], session × injury [F_(1.502, 51.05)_ = 0.91, p = 0.86, Geisser-Greenhouse corrected, ε =0.75], sex × injury [F_(1.34)_ = 0.01, p = 0.92], and session × sex × injury [F_(1.502, 51.05)_ = 1.60, p = 0.22, Geisser-Greenhouse corrected ε =0.75] (Fig. 2G). To assess magazine-checking behavior, which may reflect monitoring of the action-outcome relationship, we assessed the percent of lever presses that were followed by a magazine entry on the final day of training. A two-way ANOVA indicated no effect of sex [F_(1.31)_ = 0.19, p = 0.66], injury [F_(1,31)_ = 0.14, p = 0.71], or interaction sex × injury [F_(1,31)_ = 0.19 p = 0.66] on magazine checking (Fig. 2H).

These results indicate similar baseline behavior between SNI and sham groups, suggesting that SNI-induced hypersensitivity does not affect acquisition of operant sucrose self-administration. All groups acquired and performed the task similarly, regardless of sex or injury, with session-dependent changes reflecting normal learning throughout acquisition. Three animals (2 SNI males and 1 sham male) were excluded in this phase due to failure to acquire the task.

### SNI facilitated reversal learning in operant sucrose self-administration

Reversal learning was assessed by switching the reinforced lever so that presses the previously inactive lever now delivered sucrose, while the previously active lever had no programmed consequence. The reinforcement schedule remained VR2, with one 30-min session per day for at least 12 days.

A rmANOVA revealed a significant main effect of session on the number of presses on the newly reinforced lever [F_(3.511,114.6)_ = 52.65; p < 0.0001]. Post hoc analysis indicated that responding on the new active lever was increased after session 1 (all p’s < 0.05) (Fig. 3A). This suggests that all mice were able to acquire the reversal. There were no effects of sex [F_(1,34)_ = 0.0008; p = 0.98], injury [F_(1,34)_ = 0.03; p = 0.87], or interaction between session × sex [F_(11,359)_ = 1.52, p = 0.12], session × injury [F_(11,359)_ = 0.35, p = 0.97], sex × injury [F_(1.34)_ = 1.17, p = 0.28], and session × sex × injury [F_(11,359)_ = 0.63, p = 0.81]. In order to more closely examine any impact of SNI or sex on the initial acquisition of reversal learning, the first two sessions were analyzed. Here, a rmANOVA again showed a main effect of session [F_(1,34)_ = 12.98; p = 0.001] as well as a significant session × sex × injury interaction [F_(1,34)_ = 4.34; p = 0.05]. Post hoc tests indicated that only SNI females significantly increased lever pressing on the newly active lever on day 2 compared with day 1 (p = 0.002). In contrast, the other experimental groups did not increase lever pressing on the newly reinforced lever on day 2 compared with day 1 (p’s > 0.3). There were no effects of sex [F_(1,34)_ = 1.50; p = 0.23], injury [F_(1,34)_ = 0.85; p = 0.36], or interactions of session × sex [F_(1,34_ = 3.06, p = 0.09], session × injury [F_(1,34)_ = 0.25, p = 0.62], or sex × injury [F_(1.34)_ = 3.00, p = 0.09] (Fig. 3B).

**Figure 3.**
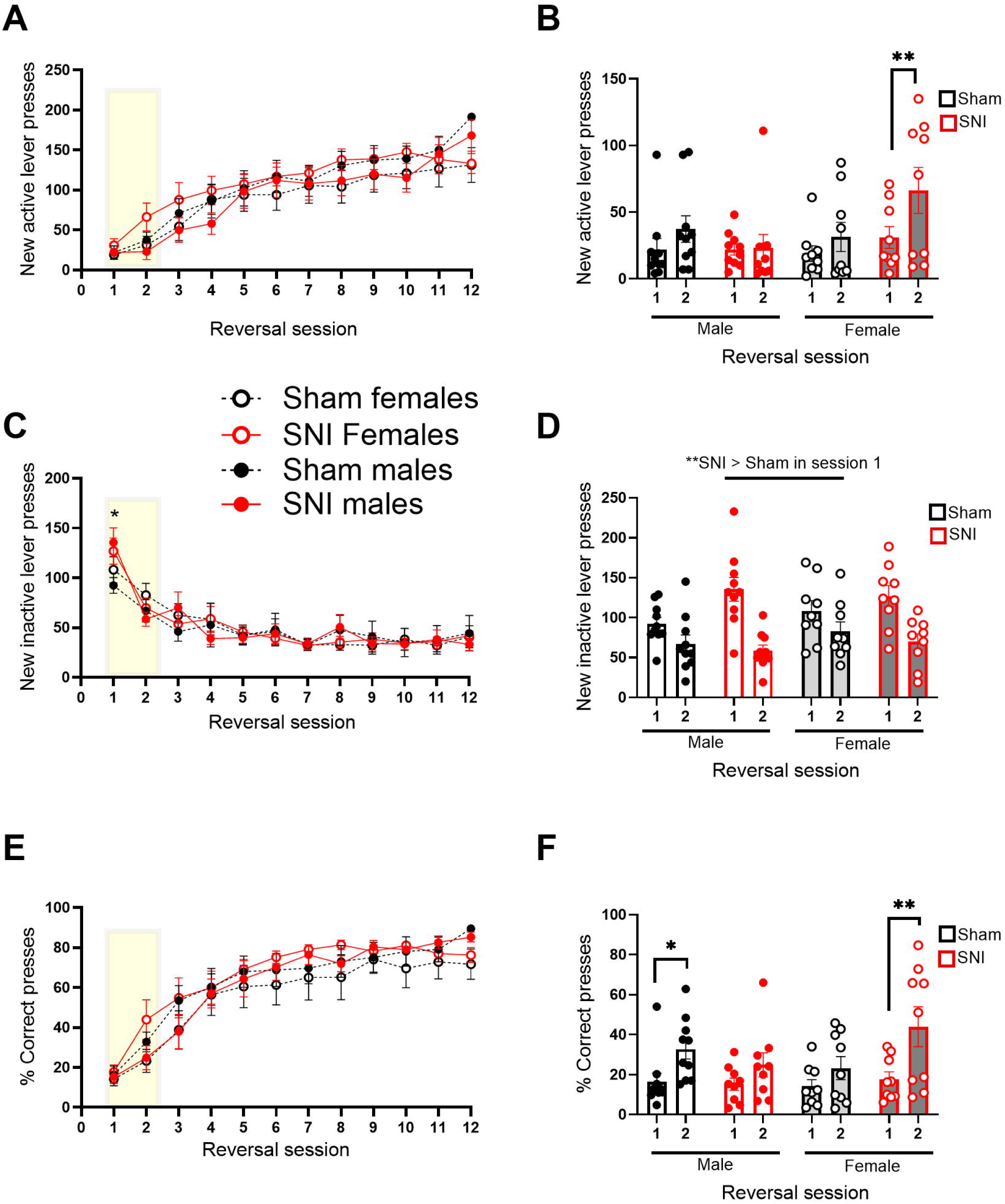
SNI facilitates initial reversal learning in operant sucrose self-administration in females. Reversal learning was assessed by switching the reinforced lever such that presses on the previously inactive lever now delivered sucrose, while presses on the previously active lever had no programmed consequence (VR2 schedule; one 30-min session per day). (A) Presses on the newly reinforced lever across reversal sessions increased over time in all groups, indicating successful acquisition of the reversal. (B) Analysis of the first two reversal sessions (yellow shadow) revealed facilitated acquisition in SNI females, which showed a significant increase in responding on the newly active lever from day 1 to day 2. No other group showed a significant increase. (C) Presses on the previously active lever across reversal sessions. SNI mice exhibited higher response than sham mice on day 1 only, indicating impaired initial inhibition of the previously reinforced response. (D) Responding to the newly inactive lever during the first two reversal sessions (yellow shadow). SNI mice showed higher responding on day 1 compared with sham mice, but both groups reduced responding by day 2. (E) The percentage of responding on the newly active lever across reversal sessions increased over time in all groups, reflecting improved response selection. (F) Analysis of the first two reversal sessions (yellow shadow) showed that only sham males and SNI females significantly increased the percentage of presses on the newly active lever from day 1 to day 2. Data are presented as mean ± SEM. n = 9–10 mice per group.

Successful reversal learning requires the inhibition of the previously reinforced behavior in addition to the acquisition of the new response. A rmANOVA revealed a significant main effect of session [F_(3.988,133.8)_ = 52.17; p < 0.0001; Greenhouse-Geisser corrected, ε = 0.36] and a significant session × injury interaction [F_(11,369)_ = 2.73; p = 0.002] on previously active lever presses (Fig. 3C). Post hoc analysis indicated that pressing on the previously active lever press was significantly higher in SNI mice compared to sham on day 1 (p = 0.015; all other sessions p’s > 0.3). There were no effects of sex [F_(1,34)_ = 0.008; p = 0.93] or injury [F_(1,34)_ = 0.009; p = 0.92], or interactions between session × sex [F_(11,369)_ = 1.34, p = 0.18], sex × injury [F_(1.34)_ = 0.02, p = 0.90], and session × sex × injury [F_(11,369)_ = 1.16, p =0.32]. For the initial phase of reversal learning, a rmANOVA revealed a main effect of session [F_(1,34)_ = 78.31; p < 0.001] and an interaction between session × injury [F_(1,34)_ = 15.80; p = 0.0003] (Fig. 3D). Post hoc comparisons indicated that the new inactive lever responding was higher on day 1 in SNI mice than in sham mice (p < 0.01) but did not differ on day 2 (p = 0.35). Both SNI and sham mice reduced responding on the new inactive lever from day 1 to 2 (p’s < 0.01). There were no effects of sex [F_(1,34)_ = 0.71; p = 0.40], injury [F_(1,34)_ = 1.00; p = 0.33], or interactions between session × sex [F_(1,34)_ = 0.90, p = 0.35], sex × injury [F_(1.34)_ = 0.51, p = 0.48], and session × sex × injury [F_(1, 34)_ = 0.89, p = 0.35].

To determine the effect of sex and injury on the percentage of “correct” responding on the new active lever out of total lever presses, the number of new active lever presses was divided by the sum of active and inactive lever presses. A rmANOVA revealed a significant main effect of session [F_(3.699,117.4)_ = 87.47; p < 0.0001, Geisser-Greenhouse corrected ε = 0.34] (Fig. 3E). Post hoc analysis indicated that the percent correct increased vs session 1 on all subsequent sessions (p’s < 0.05). There were no effect of sex [F_(1,33)_ = 0.07; p = 0.80], injury [F_(1,33)_ = 0.54; p = 0.47], or interaction session × sex [F_(11,349)_ = 0.55, p = 0.87], session × injury [F_(11,349)_ = 0.57, p = 0.85], sex × injury [F_(1.33)_ = 0.79, p = 0.38], and session × sex × injury [F_(11,349)_ = 1.60, p = 0.09]. To investigate the initial acquisition of the reversal, a rmANOVA on the first two sessions was performed. It revealed again a main effect of session [F_(1,33)_ = 37.32; p < 0.0001] and an interaction between session × sex × injury [F_(1,33)_ = 5.64; p = 0.02]. (Fig. 3F). Post hoc comparisons showed that from day 1 to day 2, only sham males (p = 0.0208) and SNI females (p < 0.001) increased the percentage of presses on the new active lever. There were no effect of sex [F_(1,33)_ = 0.26; p = 0.61], injury [F_(1,33)_ = 0.61; p = 0.44], or interaction session × sex [F_(1,33)_ = 0.84, p = 0.36], session × injury [F_(1,33)_ = 1.04, p = 0.31], sex × injury [F_(1.33)_ = 2.90, p = 0.10], and session × sex × injury [F_(1,33)_ = 5.64, p = 0.02].

Consistent with the differences observed only in the initial acquisition of the reversal, a rmANOVA of the days to reach reversal criteria did not differ by injury [F_(1,32)_ = 0.04; p = 0.84], sex [F_(1,32)_ = 0.89; p = 0.35], or injury × sex interaction [F_(1,32)_ = 1.20; p = 0.28] (data not shown).

### SNI did not impact extinction of operant sucrose self-administration

During the extinction phase, no reinforcement was delivered following presses on either lever. A REML revealed a significant main effect of session on the number of presses on the previously active lever [F_(3.74,125.9)_ = 42.10; p < 0.0001, Geisser-Greenhouse corrected ε = 0.62]. A post hoc analysis indicated that responding on the previously active lever decreased after session 1 (all p’s < 0.0001) indicating that all groups extinguished responding on this lever over time (Fig. 4A). There were no effects of sex [F_(1, 34)_ = 0.60; p = 0.45] or injury [F_(1,34)_ = 0.12; p = 0.73], or interactions between session × sex [F_(6, 202)_ = 0.31, p = 0.93], session × injury [F_(6, 202)_ = 0.58, p = 0.75], sex × injury [F_(1,34)_ = 0.06, p = 0.80], or session × sex × injury [F_(6, 202)_ = 0.69, p = 0.67]. As all mice previously underwent reversal learning, responding on the inactive lever was assessed to confirm that no resurgence in responding was observed. No differences in inactive lever pressing were observed as effects of session [F_(2.947,99.22)_ = 2.66; p = 0.053, Geisser-Greenhouse corrected ε = 0.49), injury [F_(1,34)_ = 0.54, p = 0.47] or sex [F_(1,34)_ = 1.72, p = 0.20], nor were there interactions between session × sex [F_(2.947,99.22)_ = 0.35; p = 0.79, Geisser-Greenhouse corrected ε = 0.49], session × injury [F_(2.947,99.22))_ = 0.73, p = 0.54, Geisser-Greenhouse corrected ε = 0.49], sex × injury [F_(1.34)_ = 0.004, p = 0.95], or session × sex × injury [F_(2.947,99.22))_ = 0.63, p = 0.60, Geisser-Greenhouse corrected ε = 0.49] (Fig. 4B). The number of magazine entries during extinction was analyzed by a rmANOVA, which revealed a significant main effect of session [F_(3.65,124.2)_ = 24.79, p < 0.0001, Geisser-Greenhouse corrected ε = 0.61]. A post hoc analysis indicated that magazine entries decreased after session 1 (all p’s < 0.0001) indicating that all groups extinguished this behavior over time (Fig. 4C). There were no effects of sex [F_(1,34)_ = 0.98, p = 0.31], or injury [F_(1,34)_ = 1.37, p = 0.25], or interaction between session × sex [F_(3.65,124.2)_ = 1.76, p = 0.15, Geisser-Greenhouse corrected ε = 0.61], session × injury [F_(3.65,124.2)_ = 2.49, p = 0.05, Geisser-Greenhouse corrected ε = 0.61], sex × injury [F_(1,34)_ = 0.21, p = 0.65], and session × sex × injury F_(3.65,124.2)_ = 1.05; p = 0.38, Geisser-Greenhouse corrected ε = 0.61] (Fig. 4C).

**Figure 4.**
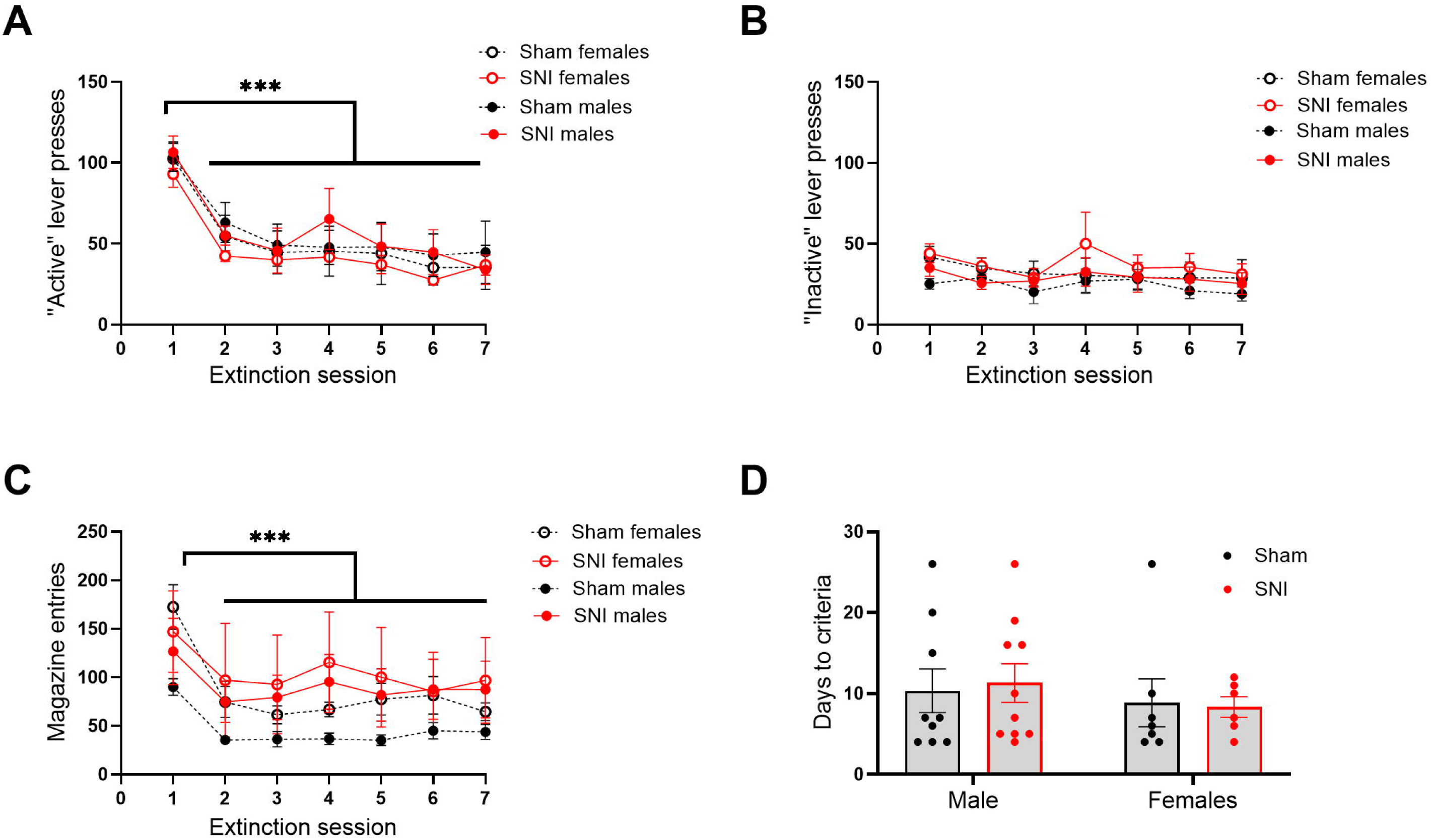
SNI does not impact extinction of operant sucrose self-administration. (A) Presses on the previously active lever decreased across extinction sessions in all groups, indicating successful extinction of sucrose-seeking behavior. (B) Presses on the previously inactive lever showed a trend toward a session effect but were not significantly affected by sex or injury. (C) Magazine entries decreased across extinction sessions in all groups, reflecting the extinction of reward-directed checking behavior. (D) The number of sessions required to reach extinction criteria did not differ by injury or sex. Data are presented as mean ± SEM. n = 9 mice per group. One sham male failed to reach extinction criteria within 30 extinction sessions and was excluded from the analysis.

A rmANOVA indicated that the number of days to reach extinction criteria did not differ by injury [F_(1, 28)_ = 0.007; p = 0.93] or sex [F_(1,28)_ = 0.73; p = 0.40], or interaction between sex × injury [F_(1, 28)_ = 0.08; p = 0.78] (Fig. 4D). One sham male failed to extinguish within 30-day sessions and was therefore excluded from further analysis.

### Painful stimulation suppressed sucrose seeking in males

To determine the effect of painful stimulation on the sucrose seeking (lever presses) and sucrose taking-related behavior (magazine entries), sham and SNI animals were stimulated 10 times with either a ‘moderate’ stimulation using the von Frey filament closest to the calculated 50% PWT, or ‘high’ stimulation using two fibers above the 50% PWT, corresponding theoretically to approximately a 90% PWT (Fig.1A). Following moderate stimulation, a REML showed no main effect of injury [F_(1, 27)_ = 0.49, p = 0.49], or session [F_(1, 24)_ = 3.90, p = 0.05] on lever pressing. However, a significant main effect of sex was observed [F_(1, 27)_ = 4.96, p = 0.03], indicating that females pressed more than males after moderate pain stimulation for the two days following moderate painful stimulation (Fig 5A). No interactions were observed between session × sex [F_(1, 24)_ = 0.14, p = 0.71], session × injury [F_(1, 27)_ = 2.24, p = 0.15], sex × injury [F_(1,27)_ = 0.38, p = 0.54] or session × sex × injury [F_(1,24)_ = 0.05, p = 0.82].

**Figure 5.**
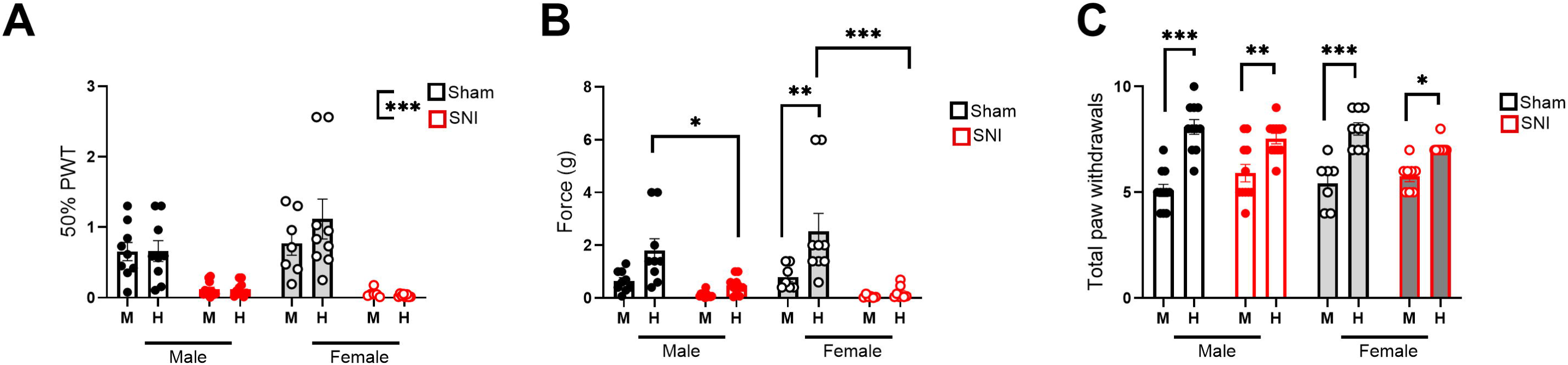
Painful stimulation suppressed sucrose seeking in a sex-dependent manner. Sham and SNI mice received 10 stimulations of either moderate (von Frey filament closest to the 50% PWT) or high painful stimulation (two filaments above the 50% PWT, corresponding to ∼90% PWT). Lever pressing and magazine entries were normalized to the average number of these behaviors during the two days preceding stimulation and assessed on the stimulation day and on the subsequent day. (A) Following moderate painful stimulation, sucrose seeking (active lever pressing) was reduced in males compared with females. SNI males and sham females exhibited suppression of lever pressing after moderate stimulation. (B) Following high painful stimulation, females again showed higher lever pressing than males, with reduced responding driven mainly by suppression of sucrose seeking in males, particularly SNI males. (C) Moderate painful stimulation increased magazine entries in a sex- and injury dependent manner, with enhanced reward magazine checking in SNI mice. (D) High painful stimulation did not significantly alter magazine entries across groups, although trends toward increased responding were observed in some male groups. The dotted gray line represents 100%. Data are presented as mean ± SEM. n = 8-9 mice per group. One SNI female was excluded due to low sensitivity to painful stimulation at the final test and removal of outliers.

To determine whether acute painful stimulation resulted in a significant change in behavior from baseline, a one-sample t test versus 100% (baseline) revealed that moderate stimulation decreased lever pressing in SNI males for two days following stimulation [one-sample t, day 1: t = 3.61, df = 9, p = 0.006; day 2: t = 6.09, df = 9, p = 0.0002] and in sham males [one-sample t, day 1: t = 2.70, df = 6, p = 0.04; day 2: t = 4.28, df = 7, p = 0.004]. Sham females exhibited suppressed responding only on the stimulation day [one-sample t, day 1: t = 3.32, df = 4, p = 0.03; day 2: t = 1.78, df = 5, p = 0.14]. SNI females did not exhibit reduced sucrose seeking following moderate painful stimulation on either day [one-sample t, day 1: t = 0.11, df = 4, p = 0.92; day 2: t = 0.61, df = 5, p = 0.57]. Together, this indicates that greater responding observed in females than males following moderate pain stimulation was driven by reduced lever pressing in males.

The same analyses were done after high pain stimulation. Analysis revealed no effects of injury [F_(1,30)_ = 3.61, p = 0.07] or session [F_(1, 30)_ = 0.89, p = 0.35], but a main effect of sex [F_(1, 30)_ = 13.83, p = 0.0008] with females pressing more than males after high stimulation for the two days following (Fig. 5B). No interactions were observed for session × sex [F_(1,30)_ = 0.006, p = 0.94], session × injury [F_(1,30)_ = 1.12, p = 0.30], sex × injury [F_(1,30)_ = 0.09, p = 0.77], or session × sex × injury [F_(1,30)_ = 2,69 p = 0.11]. To determine differences from baseline behavior, one-sample t tests versus baseline (100%) revealed that high stimulation decreased lever pressing in SNI males for two days following stimulation [one-sample t, day 1: t = 2.80, df = 9, p = 0.02; day 2: t = 3.78, df = 9, p = 0.0044]. Sham males exhibited suppressed responses only on the stimulation day and a trend on day 2 [one-sample t, day 1: t = 3.34, df = 7, p = 0.01; day 2: t = 2.22, df = 7, p = 0.06]. No significant reductions were observed following high pain stimulation in sham [one-sample t, day 1: t = 2.16, df = 8, p = 0.06; day 2: t= 0.74, df = 8, p = 0.74] or SNI females [one-sample t, day 1: t = 0.007, df = 7, p = 0.99; day 2: t = 1.24, df = 8, p = 0.99], consistent with findings that higher responding in females was driven by reductions in sucrose seeking in males following painful stimulation (Fig. 5B).

To determine the effect of painful stimulation on sucrose reward port entries, magazine entries were normalized to the average number of entries during two days preceding stimulation (Fig 5C). A rmANOVA showed a significant main effect of injury [F_(1, 28)_ = 5.77, p = 0.02] and interactions between session x injury [F_(1, 28)_ = 4.72, p = 0.04] and session x injury x sex [F_(1,28)_ = 4.32, p = 0.05]. Post hoc tests showed two days after moderate pain stimulation, SNI females exhibited greater magazine entries than sham females and males (all p’s < 0.05). No significant main effects of sex [F_(1, 28)_ = 3.05, p = 0.09] or session [F_(1, 28)_ = 0.20, p = 0.66] or between interaction session x sex [F_(1, 28)_ = 3.30, p = 0.08], sex x injury [F_(1,28)_ = 0.96, p = 0.34] or session x injury x sex [F _(1,28)_ = 4.32, p = 0.34] were observed. To determine differences from baseline behavior, a one-sample t test versus baseline (100%) revealed that moderate stimulation increased magazine entries in SNI males on day 1 and a trend on day 2 [one-sample t, day 1: t = 2.93, df = 10, p = 0.02; day 2: t = 2.00, df = 10, p = 0.07]. Sham males showed increased magazine entries only on the day 2 after stimulation [one-sample t, day 1: t = 0.89, df = 7, p = 0.40; day 2: t = 2.48, df = 7, p = 0.04]. No changes in magazine entries following moderate stimulation were observed in SNI [one-sample t, day 1: t = 1.34, df = 5, p = 0.24; day 2: t = 1.80, df = 5, p = 0.13] or sham females [one-sample t, day 1: t = 2.24, df = 6, p = 0.06; day2: t = 0;26, df = 6, p = 0.81]. Together, this suggests that moderate pain stimulation has opposing effects on magazine entries and lever presses, with modest increases in magazine entries in males (Fig. 5C).

The magazine entries after high pain stimulation were analyzed and revealed no significant effects of sex [F_(1,32)_ = 1.02, p = 0.32], injury [F_(1,32)_ = 0.02, p = 0.88], or session [F_(1,29)_ = 0.20, p = 0.66], or interactions between session x sex [F=_(1,29)_ = 0.04, p = 0.83], session x injury [F=_(1,29)_ = 0.35, p = 0.56], sex x injury [F=_(1,32)_ = 0.18, p = 0.67] or session x injury x sex [F=_(1,29)_ = 0.09, p = 0.76]. To determine differences from baseline behavior, a one-sample t test versus 100% revealed no significant changes in magazine entries following high pain stimulation in any groups [SNI males - one-sample t, day1: t = 2.07, df = 9, p = 0.07; day2: t = 0.96, df = 9, p = 0.36; sham males - one-sample t, day 1: t = 21.21, df = 7, p = 0.27; day 2: t= 1.04, df = 8, p = 0.33; sham females - one-sample t, day 1: t = 1.20, df = 6, p = 0.27; day 2: t= 2.05, df = 7, p = 0.08; SNI females - one-sample t, day 1: t = 0.874, df = 7, p = 0.41; day2: t= 0.91, df = 8, p = 0.34] (Fig. 5D). One SNI female was excluded from the analysis due to low sensitivity to painful stimulation at the final test.

### Mechanical allodynia was stable across testing

To determine if repeated stimulations would impact mechanical sensitivity, PWT was compared between “moderate” and “high” pain stimulation conditions at the same experimental timepoints described in the anterior results and shown at the timeline (Fig.1A). There were no significant effects of time [F_(1,28)_ = 1.851; p = 0.18], indicating that PWT were not different during moderate and high stimulation in the same experimental group. A main effect of injury [F_(1,33)_ = 35.04; p < 0.0001] showed a significant difference between sham and SNI, as expected, with the SNI ipsilateral paw exhibiting lower PWT than the sham ipsilateral paw in both sexes (Fig. 6A), showing that the mechanical allodynia produced by the SNI surgery was stable after 8 and 9 weeks.

**Figure 6.**
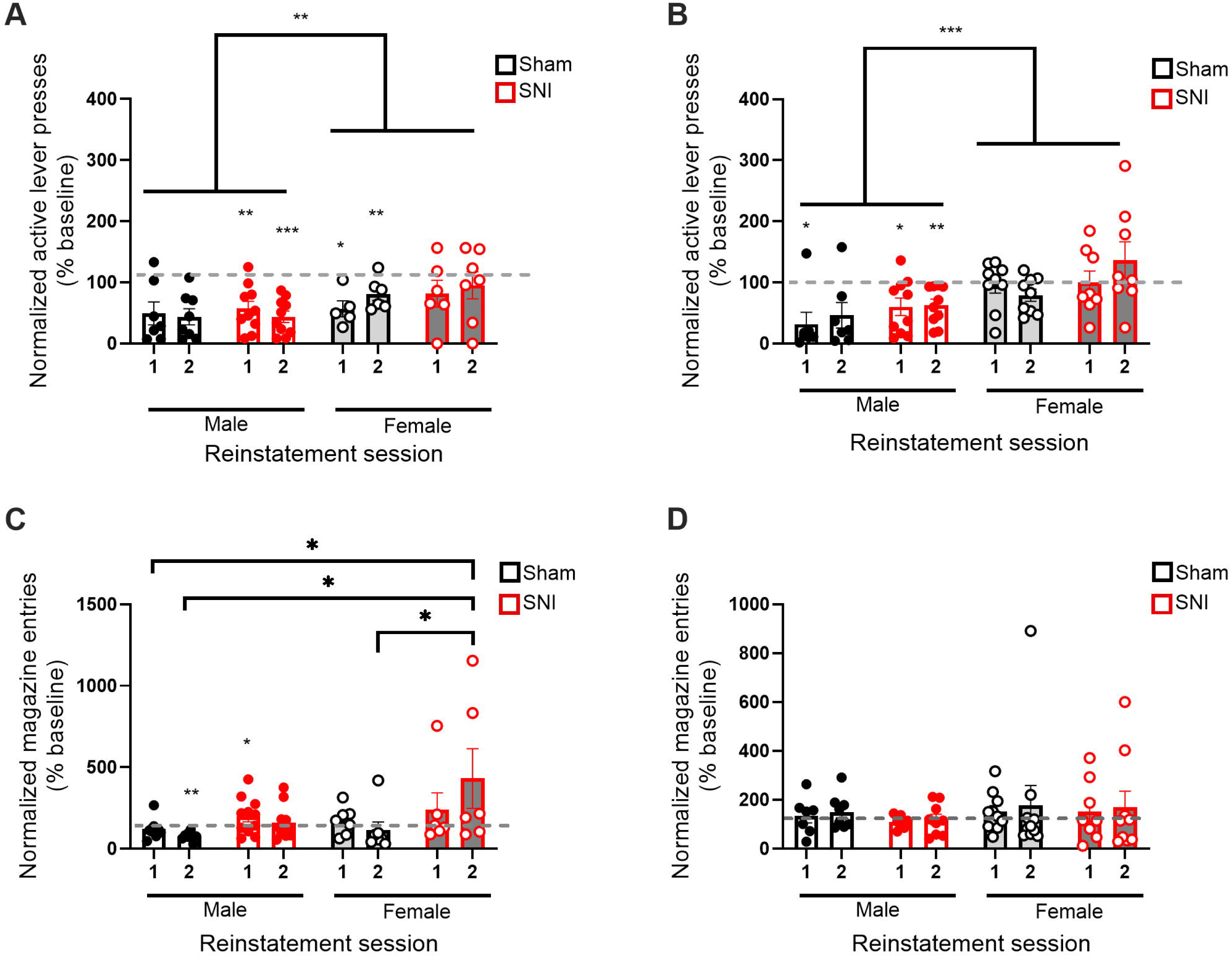
Characterization of painful stimulation in SNI and sham animals. Pain-induced reinstatement tests were separated by additional extinction sessions and counterbalanced between moderate and high stimulation days to avoid order effects. (A) Paw withdrawal thresholds (PWT) did not differ between moderate and high stimulation test days within each experimental group, indicating stable mechanical sensitivity across reinstatement sessions. As expected, SNI mice exhibited significantly lower ipsilateral PWT compared with sham mice in both sexes, demonstrating persistent mechanical allodynia 8–9 weeks after surgery. (B) Force (g) applied with von Frey filaments to the ipsilateral paw differed between moderate and high pain-reinstatement days in sham mice, but not in SNI mice, likely because the force used on the moderate stimulation day in SNI animals was already near threshold. (C) Despite differences in applied force, the number of hindpaw withdrawal responses was significantly higher during high versus moderate stimulation in all groups, indicating comparable sensory engagement across experimental conditions. Data are presented as mean ± SEM; n = 8-9 mice per group.

To compare the amount of force in grams applied with the von Frey fibers in the ipsilateral paw, a REML indicated a significant main effect of stimulation day [F_(1,28)_ = 19.76, p = 0.0001] and paw [F_(1,33)_ = 24.31, p < 0.0001] and interaction between stimulation day × paw [F_(1,28)_ = 9.951, p < 0.004]. Post hoc test shows a significant difference between the moderate and high stimulus in sham females (p = 0.0005) and sham males (p = 0.017), but not in SNI females (p = 0.99) and SNI males (p = 0.79), possibly because the force applied in the moderate pain-stimulation day was already very low in SNI, resulting in no significant difference compared with the force applied in the high pain-stimulation day (Fig. 6B).

Importantly, while the forces used to elicit paw withdrawal responses on moderate pain-stimulation day and high pain-stimulation day differed between SNI and sham animals, the number of hindpaw withdrawal responses did not differ. A REML on the number of hindpaw withdrawal responses revealed a main effect of stimulation force [F_(1, 32)_ = 110.6, p < 0.0001], and in the interaction stimulation force × injury [F_(1,32)_ = 10.02, p = 0.003]. Sidák’s multiple comparisons show a significant difference between the moderate and high stimulus in all groups, sham females (p < 0.0001), sham males (p < 0.0001), SNI females (p = 0.01), and SNI males (p = 0.0005) (Fig. 6C). One SNI female was excluded from analyses due to low sensitivity.

### Painful stimulation differentially modulates mPFC neuronal activation in sham and SNI mice

To assess the effects of painful stimulation on the activation of the mPFC (Fig. 7A) in mice with and without a history of chronic pain, c-Fos expression was analyzed by IHC in mice that underwent painful stimulation 90 minutes prior to transcardial perfusion. Mice were matched to receive either 10 stimulations with a von Frey filament corresponding to ∼ 90% PWT for their respective experimental group (shams: 2 g and SNI: 0.2 g) or serve as no stimulation controls. Because there were no differences between sexes, data from females and males were combined for the IHC analyses. A two-way ANOVA revealed no main effect of injury [F_(1,19)_ = 1.68; p = 0.21] or painful stimulation [F_(1,19)_ = 0.43; p = 0.52], or interaction between pain stimulation x injury [F_(1,19)_ = 3.51; p = 0.08] in PrL (Fig. 7B). In the IfL, there was no main effect of injury [F_(1,20)_ = 3.05; p = 0.10] or painful stimulation [F_(1,20)_ = 0.62; p = 0.44] but there was a significant interaction between pain stimulation x injury [F_(1,20)_ = 14.11; p = 0.001]. Post hoc analyses showed that SNI-stimulated animals had higher c-Fos expression in IfL compared with SNI non-stimulated animals (p = 0.006) and sham-stimulated animals (p = 0.0006). In contrast, sham mice that underwent painful stimulation exhibited lower IfL c-Fos expression compared with sham animals not stimulated (p = 0.04) (Fig. 7C). In the ACC, there was no main effect of injury [F_(1,22)_ = 0.88; p = 0.36] or interaction between injury x pain stimulation [F_(1,22)_ = 0.25; p = 0.62], but there was a main effect of pain stimulation [F_(1,22)_ = 5.39; p = 0.03] (Fig. 7D). Post hoc indicated that painful stimulation decreased c-Fos expression on ACC in sham animals only (p = 0.04) whereas SNI did not show significant difference. Overall, these results suggest that painful stimulation differentially modulated mPFC activity depending on injury status, increasing c-Fos expression in the IfL of SNI mice while reducing IfL and ACC activation in sham mice, with no effects observed in the PrL.

**Figure 7.**
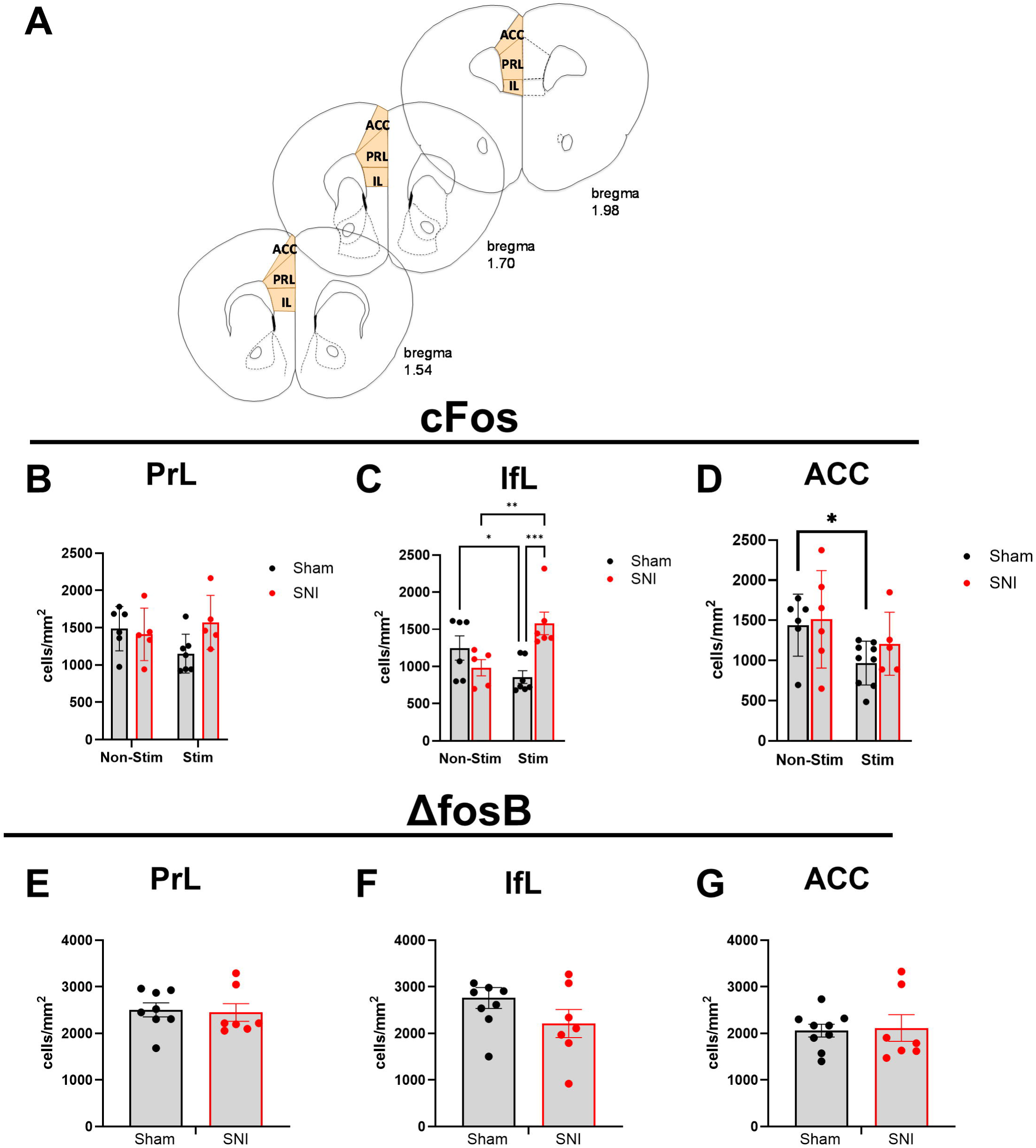
Painful stimulation differentially modulated mPFC neuronal activation in sham and SNI mice. To assess the effects of acute painful stimulation on medial prefrontal cortex (mPFC) activation in mice with or without a history of chronic pain, c-Fos expression was analyzed by immunohistochemistry. Mice received either 10 stimulations of a von Frey filament corresponding to ∼ 90% paw withdrawal threshold (PWT) for their experimental group (sham: 2 g; SNI: 0.2 g) or served as non-stimulated controls. As no sex differences were observed, data from males and females were combined. (B) In the prelimbic cortex (PrL), painful stimulation did not significantly alter c-Fos expression and did not differ by injury status. (C) In the infralimbic cortex (IfL), painful stimulation exerted opposite effects depending on injury status: c-Fos expression increased in SNI mice following stimulation, whereas sham mice exhibited reduced c-Fos expression after stimulation. (D) In the anterior cingulate cortex (ACC), painful stimulation reduced c-Fos expression in sham mice, with no significant effect observed in SNI mice. (E–G) Expression of ΔFosB, a marker of chronic neural activation, did not differ between sham and SNI mice in the PrL, IfL, or ACC. Data are presented as mean ± SEM.

To analyze the effect of chronic pain on chronic mPFC activation, ΔFosB - a marker of chronic neural activity - expression was assessed. Unpaired t-test with Welch’s correction revealed no significant difference between sham and SNI groups in PrL [t = 0.24; df = 11.9; p = 0.82] (Fig. 7E), IfL [t = 1.46; df = 11.83; p = 0.17] (Fig. 7F), or ACC [t = 0.18; df = 8.68; p = 0.86] (Fig. 7G) suggesting that this metric of chronic activation was not impacted by SNI.

## Discussion

The present study examined how chronic neuropathic pain altered reward-related behavior across a range of tasks with different cognitive and behavioral demands. SNI produced stable mechanical allodynia without disrupting acquisition or extinction of sucrose self-administration. However, chronic pain selectively facilitated reversal learning in SNI females. Sex determined the behavioral consequences of acute pain, revealing dissociable effects on lever pressing and magazine entries in males, independent of SNI. Painful stimulation differentially modulated mPFC activity depending on injury status, increasing c-Fos expression in the IfL of SNI mice while reducing IfL and ACC activation in sham mice, with no effects observed in the PrL. These findings indicate that chronic pain did not modify generally impact behavior but selectively promoted reversal learning in females, while sparing extinction learning, consistent with differential effects of chronic pain specific behavioral components.

As expected, SNI induced long-lasting mechanical hypersensitivity in the injured paw that persisted throughout behavioral testing [7,20,52,64]. This did not impair acquisition of sucrose self-administration. This aligns with studies indicating that moderate motivational demands in short-access operant tasks (e.g., 30-min sessions) may be insufficient to reveal pain-related deficits in motivation or effort [51], and with recent evidence from intravenous self-administration studies showing that chronic neuropathic pain does not impair acquisition of reward-seeking behavior but instead alters within-session acquisition dynamics and motivation under higher effort demands, particularly for drug rewards [41]. Chronic pain is increasingly recognized as a condition that alters complex learning and cognition [11,44]. For example, rats with chronic neuropathic pain retain the ability to acquire basic reward contingencies but show selective impairment in cognitive flexibility when contingencies changed, favoring previously learned associations. These deficits were accompanied by a more complex behavioral pattern characterized by burst-like responding followed by pauses, suggesting the emergence of an adaptive or compensatory learning strategy rather than a generalized learning deficit [17]. Together, these findings support the view that chronic pain promotes a shifts response strategies while largely maintaining performance.

In these experiments, female mice with chronic neuropathic pain exhibited faster acquisition of reversal learning. This pattern suggests that, in females, chronic neuropathic pain may facilitate updating of action–outcome associations. Despite this apparent enhancement in reversal learning, chronic neuropathic pain selectively altered inhibitory control during the early phase of reversal. These findings are consistent with previous evidence demonstrating that chronic neuropathic pain is accompanied by anxiety- and depressive-like behaviors in both sexes, while male mice are more vulnerable than females to pain-associated cognitive deficits [70]. Although all groups ultimately acquired reversal, SNI mice as a group showed impaired suppression of responding on the previously reinforced lever during initial reversal. This dissociation indicates that neuropathic pain did not uniformly enhance behavioral flexibility, but rather differentially affects distinct components of reversal learning, including contingency updating and inhibitory control. Consistent with this framework, chronic pain has been shown to impair cognitive flexibility and bias behavior toward perseverative or suboptimal strategies during contingency shifts, despite intact acquisition of initial action–outcome associations [17]. Together, these findings highlight that the effects of neuropathic pain on behavior interact with sex and task demands, rather than reflecting a global facilitation or impairment of cognitive flexibility.

The effects of acute pain on reward-related behaviors were largely sex specific. Both moderate and high pain stimulation suppressed sucrose seeking in extinction, as indexed by lever pressing, in male mice. Female mice persisted in sucrose seeking behavior following painful stimulation, independently of the chronic pain. In contrast to lever pressing, magazine entries were not suppressed following painful stimulation, and in fact increased in some groups following moderate pain stimulation. While these tests were conducted in extinction – and thus no reward was available/consumed – this suggests a dissociation between seeking behaviors (lever pressing) and approach or consummatory behaviors (magazine entry) [43]. We and others have observed dissociation of these factors in a range of behavioral assays, including incubation of reward seeking over abstinence [48] and habitual reward seeking [35]. We recently reported that acute painful stimulation promoted a reinstatement of ethanol seeking in a conditioned place preference paradigm in male mice [52]. The divergence observed here, where painful stimulation reduced reward seeking in males, may reflect a range of factors. One primary difference may be the nondrug reward/reinforcer used here. Ethanol is known to have analgesic properties [53,65,72] and thus may have been sought for its negative reinforcing analgesic effects. This is potentially consistent with dissociable effects of chronic pain on opioid versus sucrose reward [41]. Alternatively, this may reflect differences between the conditioned place of preference and operant self-administration paradigms. Indeed, it is possible that the reinstatement of place preference parallels the increased magazine entries/reward checking behavior observed following moderate pain stimulation. In either case, these findings suggest that acute pain effects on sucrose-reward seeking in extinction were largely independent of a history of chronic pain. It is not clear how behavior would have been impacted if tests were not conducted in extinction as aversive experience can generally suppress sucrose consumption.

At the neural level, we observed increased c-Fos expression in the infralimbic cortex of SNI mice following acute pain stimulation. The IfL is thought to facilitate the encoding and updating of contingencies between cues and behaviors to guide adaptive responding [10,34,50,55] and is required for learning alternatives to prelimbic-promoted associations through reciprocal connectivity within the medial prefrontal cortex [47]. Moreover, converging evidence indicates that IfL-centered circuits promote habitual and inflexible behavioral strategies [9]. In this context, the increased recruitment of IfL neurons in SNI mice may reflect pain-induced reweighting of prefrontal control strategies, biasing behavior toward inflexible or habit-like responding under aversive conditions. This is consistent with the hypothesis that IfL functional reorganization contributes to regulation of neuropathic pain and the modulation of reward-related behaviors [46]. Conversely, artificial electrical or pharmacological activation of the IfL decreased pain sensitivity in animals with chronic pain, further supporting a central role for IfL in integrating nociceptive and motivational signals.

In contrast, the reduction in c-Fos expression observed in the ACC following painful stimulation in sham animals was unexpected, given that both experimental and clinical studies typically report increased ACC activation in response to pain [3,25,62]. Indeed, meta-analyses of experimental pain consistently identify the ACC as a core component of the pain-processing network, alongside somatosensory, insular, prefrontal cortices, and thalamus [3,53,54]. One possible explanation for this discrepancy relates to the anatomical level analyzed in the present study. Here, ACC activity was quantified at a relatively rostral level (approximately +1 to +2 mm from bregma), whereas pain-related ACC activation is more commonly reported in more caudal subregions. Previous work has demonstrated functional heterogeneity along the rostrocaudal axis of the ACC, with caudal ACC subregions being more strongly engaged in affective and nociceptive processing, including during chronic stress and neuropathic pain states. Thus, the observed decrease in c-Fos expression may reflect region-specific modulation rather than a global suppression of ACC involvement in pain. Excitatory neurons in the ACC dynamically signal performance monitoring and rule updating during reversal learning, supporting effects on cognitive flexibility [73] but not for simple reward-driven actions [2]. This may be driven by ACC theta oscillations during reversal learning [71]. Together, the ACC is implicated in reversal learning, and rostrocaudal differences within the ACC should be considered in future analyses.

Chronic pain has been shown to alter motivational processes in a reward- and sex-dependent manner. Previous work demonstrated that male, but not female, mice exhibit increased within-session acquisition rates yet display a reduced willingness to work for opioid reward under higher effort requirements, as evidenced by fewer reinforcers earned and a lower breakpoint in progressive ratio schedules [42]. This framework is consistent with the selective effects of chronic pain observed in the present study, in which alterations emerged during early reversal and under increased demands, rather than as a general deficit in learning.

The prefrontal cortex plays a central role in inhibitory learning and behavioral regulation during addiction and relapse-related processes [19,26,29,33,55]. We observed no differences in ΔFosB expression across mPFC subregions, indicating that chronic neuropathic pain did not induce sustained baseline alterations in prefrontal activity, but instead alters circuit responsiveness to acute painful events. This is a blunt metric, however, and should be interpreted in the context of electrophysiological and functional imaging data.

One caveat to the current studies is that all animals underwent reversal learning prior to extinction training and test sessions. This prior reversal learning may facilitate extinction learning and/or result in less stable action–outcome associations, making lever pressing less stable. This is potentially consistent with increased magazine entries following moderate painful stimulation, as the reward port was consistently associated with reward delivery prior to extinction.

Overall, these findings demonstrate that chronic neuropathic pain selectively modulates cognitive and reward-related behaviors in mice. While baseline self-administration remained intact, chronic neuropathic pain facilitated reversal learning in female mice. In contrast, acute pain suppressed reward seeking in male mice, independent of chronic pain status. Behavioral alterations were accompanied by injury-dependent recruitment of IfL and ACC, emphasizing the mPFC as a critical site for the interaction between pain and reward-related behavior.

## Acknowledgements

This research was supported by the National Institute on Alcohol Abuse and Alcoholism under Award Number R21AA027629 (JMB) and São Paulo Research Foundation (FAPESP), grant 2024/09394-6 (AAB). Data will be made available at Open Science Framework.

## Author Contributions

Conceptualization – JMB, AAB, MAN Data curation – AAB

Formal analysis – AAB, JMB Funding acquisition – AAB, JMB Investigation – AA, MAN, CMCA

Methodology – AAB, MAN, JMB, CMCA Project administration – AAB, JMB Supervision - JMB

Visualization – AAB, JMB

Writing – original draft – AAB, JMB

Writing – review and editing – AAB, JMB, KPA, MAN, CMCA

